# Lattice-based microenvironmental uncertainty driven phenotypic decision-making: a comparison with Notch-Delta-Jagged signaling

**DOI:** 10.1101/2021.11.16.468748

**Authors:** Aditi Ajith Pujar, Arnab Barua, Divyoj Singh, Ushasi Roy, Mohit Kumar Jolly, Haralampos Hatzikirou

## Abstract

Phenotypic decision-making is a process of determining important phenotypes in accordance with the available microenvironmental information. Although phenotypic decision at the level of a single cell has been precisely studied, but the knowledge is still imperceptible at the multicellular level. How cells sense their environment and adapt? How single cells change their phenotype in a multicellular complex environment (without knowing the interactions among the cells), is still a rheotorical question. To unravel the fragmental story of multicellular decision-making, Least microEnvironmental Uncertainty Principle (LEUP) was refined and applied in this context. To address this set of questions, we use variational principle to grasp the role of sensitivity, build a LEUP driven agent-based model on a lattice which solely hinges on microenvironmental information and investigate the parallels in a well-known biological system, viz., Notch-Delta-Jagged signaling pathway. The analyses of this model led us to interesting spatiotemporal patterns in a population of cells, responsive to the sensitivity parameter and the radius of interaction. This resembles the tissue-level pattern of a population of cells interacting via Notch-Delta-Jagged signaling pathway in some parameter regimes.

## 1 Introduction

Decision-making is an action to find favorable alternatives based on certain goals, which is further shaped by multiple factors ^1^. Similarly, decision-making can be defined on the cellular levels. At first, chemicals/nutrients diffuse to the surface of the cell, which later bind to the receptors and generate a set of biochemical reactions inside the cell ^2^. After that, the cell gives feedback to their local microenvironment in terms of migration, differentiation, division, or apoptosis. The key ingredients of cell decision-making are (**I**) internal components (e.g., metabolites concentration, genetic network etc.), (**II**) external/microenvironmental components (e.g., ligands, density of cells etc.) and (**III**) the noise components which can be intracellular or intercellular ^3^. Single cell decision-making has been examined meticulously since a long time ^4^, but the study of cell decision-making on the multicellular level is obscured.

To moderately answer the question, a principle known as Least microEnvironmental Uncertainty Principle (LEUP) was developed ^2^. The principle is based on statistical mechanics tools, which was practiced in different biological setups ^5–7^. It adopts the idea of Bayesian inference ^8^ to construct the mathematical framework. The main assertion of this principle is to modify the dynamics of internal networks corresponding to the microenvironmental components, which further helps the cell to take better decisions. Although we have vast information about the molecular markers, but the knowledge about the internal dynamics which associate an internal timescale is pretty much unknown. As the number of molecular markers are immense, the inference through Bayesian formalism is obstructed. To prevent the problem, LEUP uses the notion of the minimization of microenvironmental uncertainty over time, which later helps the cell to develop the tissue. The inspiration behind this principle comes from Friston’s free energy principle ^9^ and Bayesian Brain hypothesis ^10^. Despite the fact that analogous concepts can be seen in the significant work of W.Bialek ^11^, to figure out the performance of the phenotypic dynamics driven by the LEUP theory, we compared our model with the mechanistic Notch-Delta-Jagged model ^12,13^ for a particular set of parameters. We found a good agreement between the patterns of entropy-driven model and the mechanistic model of Notch-delta-Jagged signaling. Furthermore, we have compared the order parameters in both models. Notch-delta-Jagged signaling is one of the most widely studied cell signaling mechanisms due to its versatility and simplicity ^14^. It is the heart of Epithelial-Mesenchymal transitions, cancer progressions and cell-fate determinations. Epithelial cells are stationary, and we can observe the apico-basal polarity. But, on the other hand, Mesenchymal cells are way different in nature than the epithelial one. Mesenchymal cells are non-stationary, the shape of the cell is long and which contain spherical nucleus with a prominent nucleolus. Epithelial-Mesenchymal transition is a phenomenon where Epithelial cells shift their epithelial phenotype (i.e., cell–cell adhesion and apico-basal polarity) and converge to a Mesenchymal phenotype (i.e., migratory and invasive traits) ^15^. Thus, the Notch-Delta-Jagged pathway is crucial in various biological processes including development, cancer progression, etc. Notch, a receptor on the cell membrane, binds to Delta and Jagged (ligands) from other cells. This binding helps in communicating with other cells, and the level of these proteins in a cell determines the state of the cell. Recent mechanistic models have been able to explain the patterning and its disruption in different scenarios.

Another interesting point of cell decision-making is to understand how a cell controls their sensitivity according to complex microenvironments. To gain knowledge about the cellular environment, cells, need good sensors i.e., the receptors. There exists many kinds of mechanisms like the binding of diffusible ligands ^16^, pseudopodia extension ^17^, mechanosensing ^18^, proton-pump channels ^19^, gap junctions, etc. by which cells perceive their environments. Now the question becomes, how can we quantify the phenotypic distribution of a cell in a multicellular environment without knowing the interaction among the cells in the neighborhood? How does sensitivity play a role in cellular adaptation? Can we quantify the magnitude of sensitivity in terms of a known biophysical process of internal and external variables?

To answer this question, at first, the relation between the sensitivity parameter and the dynamics of internal and external variables is derived, which was further used in a model of biophysical dynamics of a Notch-Delta-Jagged signaling ^12,13^. Furthermore, we have shown how adaptation of a cell depends on the ratio of biophysical forces and the variance of the microenvironmental probability distribution.

We have arranged this article in the following way: in Section (2) we have reviewed the Bayesian decision-making and the derivation of the phenotypic Langevin equation. Furthermore, we have calculated the sensitivity parameter using a small noise approximation to apply in Notch-Delta-Jagged model. After that, in Section (3) we studied the pattern formation and phase transition regimes for different values of sensitivity i.e., *β* and interaction radius i.e., *r* in different kinds of lattices. Finally, we discuss and conclude the outcomes of our analyses in Section. (4)

## 2 Mathematical framework

### 2.1 Bayesian inference for cell decision-making

The cellular decision-making is based on two parts i.e., (a) internal variables (i.e. representing genes, RNA molecules, translational proteins, metabolites, receptors, phenotypes etc.) and the (2) external variables (i.e. ligands, chemicals, nutrients, cellular density or stress fields e tc.). The local information sensed by the cell concealed in the internal variables. The external factors of *n*-th cell are defined as 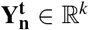 and internal factors of *n*-th cell are denoted by 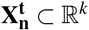, where the variables are *k*-dimensional. From a cellular Bayes an decision maker perspective, the posterior distribution 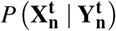 at time *t*, can be expressed as

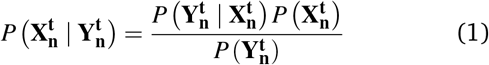

Here, the local microenvironment knowledge gaine cell, i.e., the likelihood function is shown as 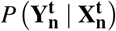 This likelihood function is modified by a multiplication term of the ratio of the probability distribution of internal variables, 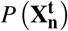., the prior and external variables 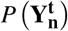. The cost of recruiting receptors to the surface of the cell encompassed an energy which is controlled further by the likelihood term, i.e., 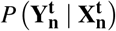. A biological example can be the response of the cell in a nutrient enriched environment. Cells sense their local environment through different biochemical performances, like polymerizing pseudopodia, translocating receptor molecules or modifying its cytoskeleton according to mechanical signals ^20,21^. If cells build an informative prior, then the price for recruitment can be reduced over time *t*.

As the information of the microenvironment is coupled with the information of internal state, the adaptation procedure of cellular internal state simultaneously enhanced with the local microenvironment, which further leads to the idea of the maximization of the internal entropy with respect to the microenvironmental entropy which behaves like a goal function to a corresponding tissue which is enclosed by a bigger set of phenotypes and the normalization of internal components. The evaluation of internal states at the steady state in terms of variational principle can be written as

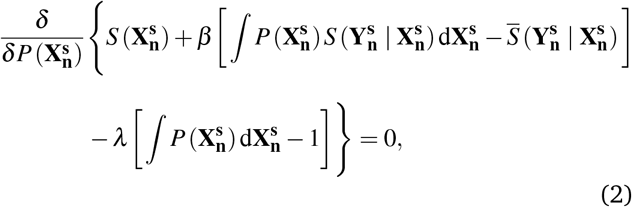

The functional derivative with respect to the internal variables is written as 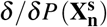. The constraints are associated with the Lagrange multipliers in Eq. (2), i.e., *β* and *λ* which corresponds to the steady state value of the local microenvironmental entropy 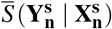 and the normalization constant of the probability distribution of the internal states. The extra information of internal and external variables from biological experiments can be written in terms of statistical observable as additional constraints. Solving Eq. (2), the probability distribution of internal states which looks like *Boltzmann*-type, i.e.,

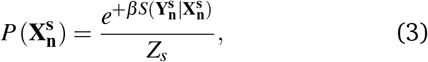

where the normalization constant is 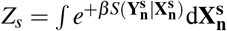 · *β* parameter leads to the sensitivity of the microenvironment. From Eq.(3), we can see that the quantity 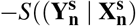 is akin to the potential of the system. In the next section, we shall use this analogy to further write the phenotypic Langevin equation for the *n*-th cell.

### 2.2 Phenotypic Langevin equation

During cellular decision-making, a cell changes its phenotype according to the states of its neighboring cells. Through Langevin equation, one can describe the phenotypic dynamics of the cell, interacting with their neighborhood. We assume that the phenotype of the *n*^th^ cell at time *t* is a continuous variable, i.e., 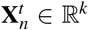. The phenotype of other cells on a two-dimensional lattice inside the neighborhood, i.e. within a distance of 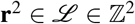 at time *t* is denoted by 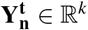 as shown in Fig. (1). As the phenotypic expression of the *n*^th^ cell changes slowly, one can consider an overdamped Langevin equation for the phenotypic time evolution. We consider that the rate of change of the phenotype is associated with the gradient of microenvironmental entropy and Gaussian noise (present during the decision-making). So, the noise has a un t variance and zero mean. The properties of the noise are 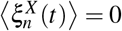 and 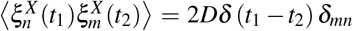 where *t*_1_ and *t*_2_ are the two distinct time points. The diffusion constant in the space of phenotype is denoted by *D*. The Delta function is defined by *δ* (*t*_1_ − *t*_2_) and the Kronecker delta defined by *δ*_*mn*_. Recalling the microenvironmental entropy being analogous to apotential, the Phenotypic Langevin’s equation for the *n*-th cell reads

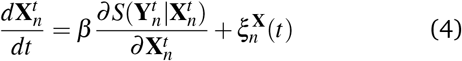

**Fig. 1.**
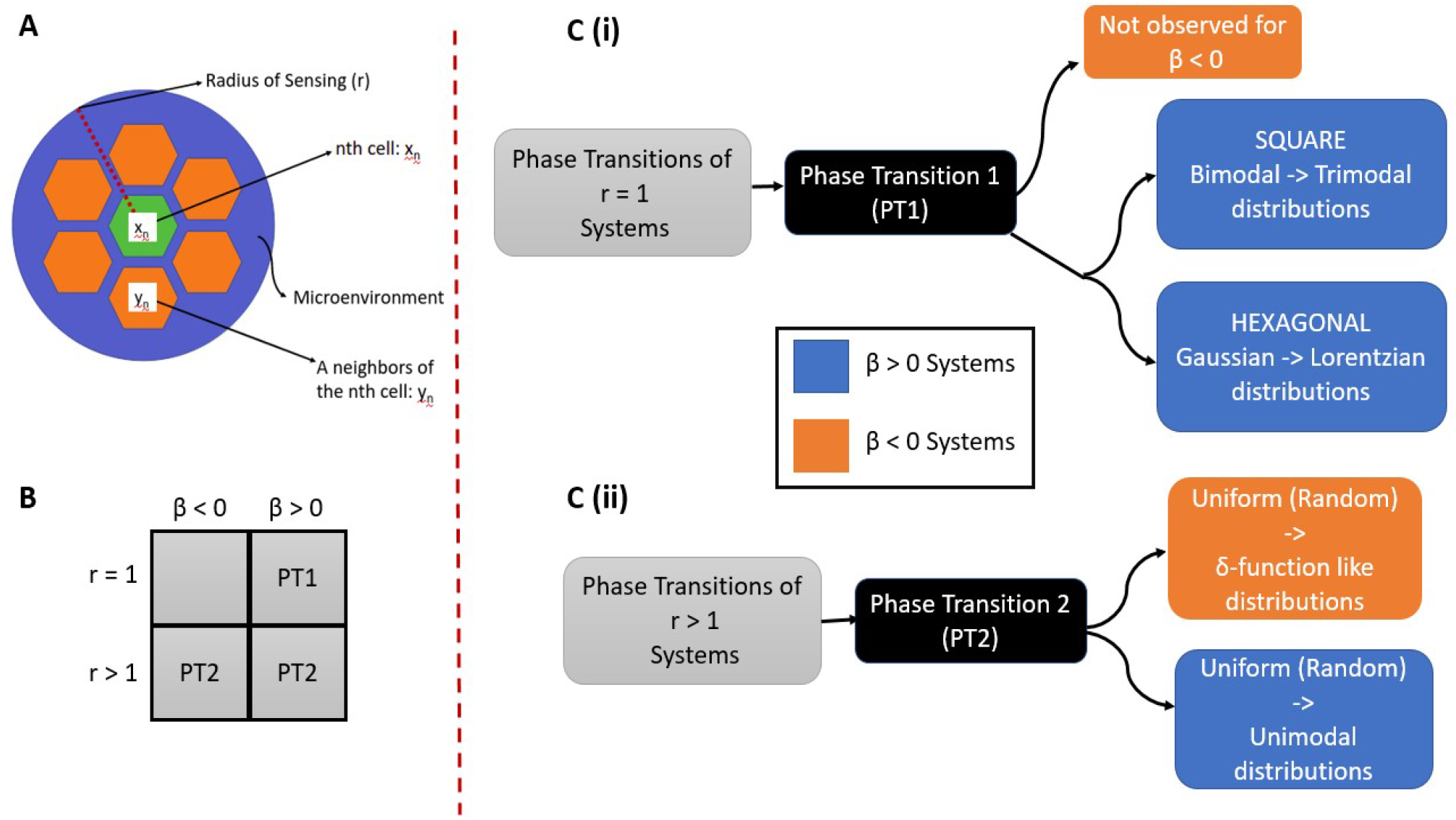
(**A**) A schematic showing internal and external variables. (**B**) Four broad parameter regimes of (±*β,r* ≥ 1) for different phase transitions. (**C**) Changes in each phase transition type with increase in |*β*|. **(i)** *PT*1 observed only for *β* > 0, where population distributions of phenotype change (differently for square and hexagonal lattices) **(ii)** *PT*2 is observed for both, positive and negative *β* values, and is similar across square and hexagonal lattices.

At a small noise approximation, one can consider the above Eq. (4) for *n*^th^ cell in a population with phenotype 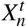 evolves according to the following equation:

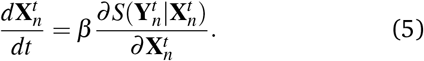

Furthermore, we have assumed that each cell senses its microenvironmental phenotypes from a *Gaussian* distribution. Further assuming that the phenotypes are all scalars (the number of dimensions of the internal variables, *k* = 1), one can construct the microenvironmental entropy for the phenotypes of *N* neighborhood cells 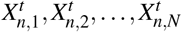 (these constitute the 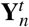) as

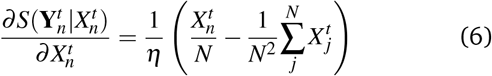

where,

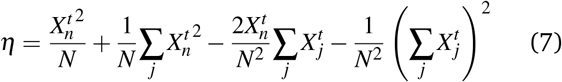

In this mathematical model, the key parameters are (**1**) the sensitivity parameter *β* and (**2**) the microenvironmental radius of interaction *r*. The latter defines the size of the sensing neighborhood, i.e., all the cells within a distance *r* from a cell will contribute to the RHS of (6) and influence the cell’s decision-making. Thus, the two parameters can help us to understand the full range of collective phenotypic behavior.

### 2.3 Mathematical model of Notch-Delta-Jagged signaling

Notch, which is a transmembrane receptor, binds to the ligands Delta and Jagged of the neighboring cell. This trans (with other cells) interaction results in the cleavage of the NICD (Notch Intra-cellular domain). NICD then goes to the nucleus and indirectly modulates the production of all 3 proteins. It transcriptionally activates Jagged and Notch and represses Delta. This asymmetric regulation results in different types of patterns in Delta dominated vs Jagged dominated system. *N*_0_, *D*_0_, *J*_0_ represent the basal production rates of the proteins. The functions *H*^*s*+^/*H*^*s*−^ represent the transcriptional activation and inhibition of the corresponding species due to NICD. *k*_*C*_ corresponds to the cis-inhibition rate. Here, the Notch binds to the Delta or Jagged of the same cell and forms a complex and hence removed from the system. *k*_*T*_ represents the transactivation rate which corresponds to the interaction with the neighboring cell’s Notch, Delta or Jagged. In the last terms, *γ* represents the basal degradation rate of the species. We assume first order kinetics for this.

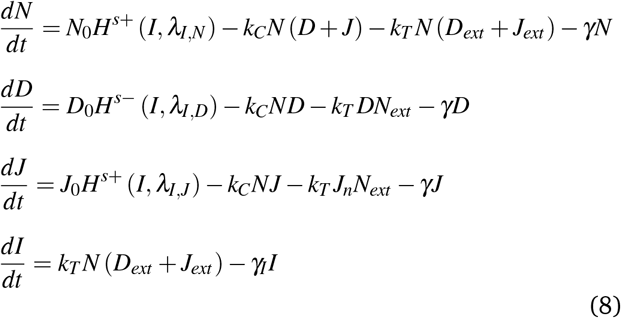

### 2.4 The relationship between sensitivity (*β*) and the internal-external coupled dynamics

The sensitivity parameter *β* quantifies the cell’s ability to sense and integrate information about their microenvironment. It can also tell us how a cell will adapt to the local microenvironment. Here we focus on the question of calculating *β* when the sensing biophysical processes are known and well described by a set of differential equations, as, for instance, in the case of ligand-receptor dynamics. To answer this question, for the *n* th cell, we consider the (*k* − dimensional) external variables 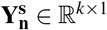, such as ligand densities, and internal variables, 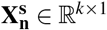, such as receptors or internalized proteins. In the following, we assume the combined dynamics of internal and external variables in the steady state:

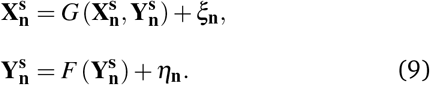

Here, we assume that the aforementioned noises follow the following multivariate Gaussian distributions 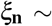 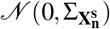 and 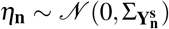. The probability distribu-tion of internal variables according to the aforementioned dynamics reads as

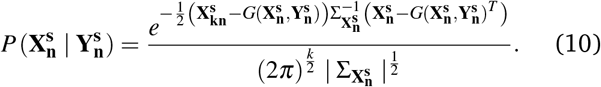

Using the small-noise approximation, and arguments as shown in ^11^, we can calculate the connection between the internal and external variable probability distributions (akin to a change of variables during integration)

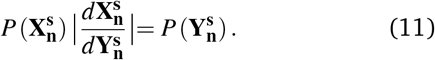

The quantity 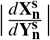 corresponds to the determinant of the Jacobian 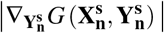 estimated at the steady state. Furthermore, we substitute the value of Jacobian at small-noise approximation limit and the value of 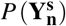 Eq. (11):

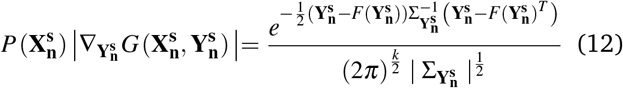

At this point we use the LEUP steady state as calculated in Eq. (3). Entropy of a *k*-dimensional multivariate normal distribution is 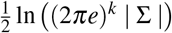. Thus,

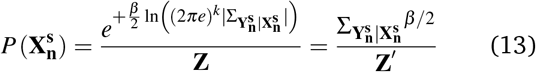

where the normalisation constants are 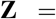 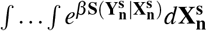 and 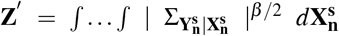 Substituting into (12), we get

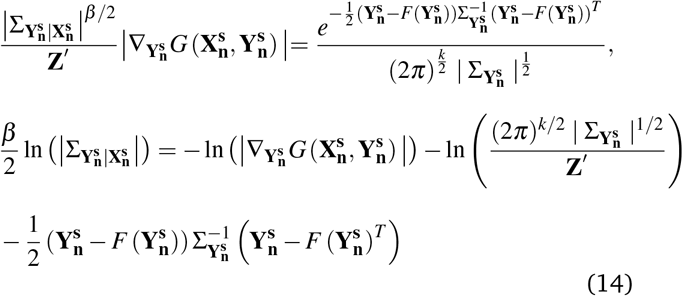

Now, the quadratic term related to the external dynamics Gaussian is assumed to be zero, by assuming that the random variable 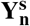 is always very close to its expected value 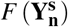. The next challenge is to come up with an estimate for the partition function. We obtain a relation for 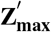 (Please check S.I. for the derivation). Taking 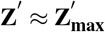, we get

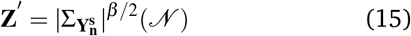

where 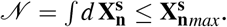. There are biological limits on the number of receptor biomolecules, that quantify the estimate for 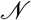. For large 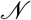, we make the simplifying assumption that ln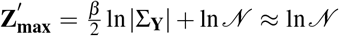. Upon substituting into Eq. (14), we then obtain

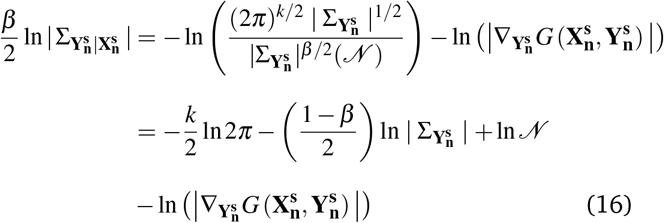

We can write finally write Eq.(16) in a compact analytic form as

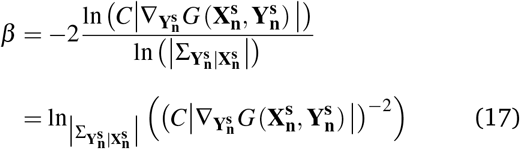

where 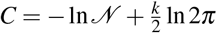.

In the next subsection, we use an example of Notch-Delta-Jagged signaling to quantify the value of sensitivity.

#### 2.4.1 Sensitivity (*β*) calculation for Notch-Delta-Jagged signaling

In this section, we shall consider a system of equations of Notch-Delta-Jagged signaling ^12,13^ with a presence of Gaussian noise. Let’s consider the Notch expression of *n*-th cell is defined by 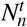, the Delta expression of *n*-th cell is defined by, 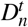 and the Jagged expression of *n*-th cell is defined by 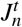. The NICD expression of *n*-th cell is defined by 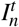. Using these sets of notations, we can write the system of equations for Notch-Delta-Jagged signaling as

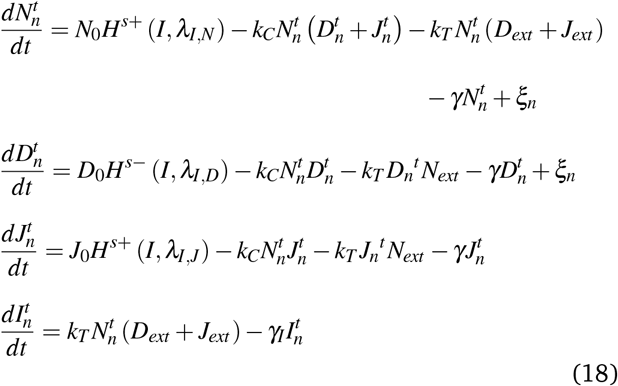

Please note that, in the mean field limit, the system of equations i.e, Eq. (18) corresponds to system of equations Eq. (8), which is the mechanistic mathematical model of Notch-Delta-Jagged signaling^12,13^. The constants *N*_0_ and *D*_0_ are the production rates of Notch and Delta proteins, which is measured in the order of molecules/h. On the other hand, *I*_0_ is the minimum value of the NICD complex Hill function in the order of molecule ^12,13^. The dissociation constant of the shifted Hill function is, *I*_0_ and the Hill coefficient is defined *h*. In the system of Eq. (18) if we add the last two equations, i.e., the Delta and Jagged equations then one can write as

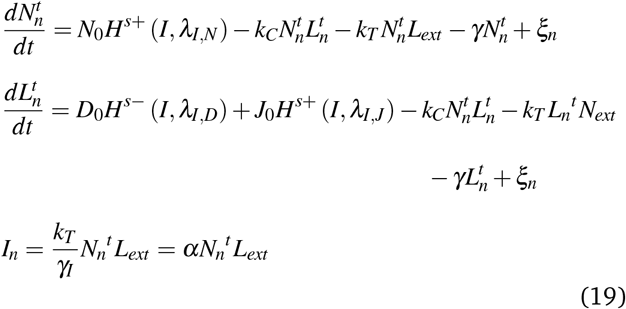

Here we define 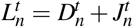 and *L*_*ext*_ = *D*_*ext*_ + *J*_*ext*_. The parameter *α* is 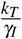. Using the quasi steady state of the NICD complex, we can substitute the value of 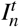 in Eq. (19). Now, one can get a simplified equation for Notch-Delta-Jagged signaling as

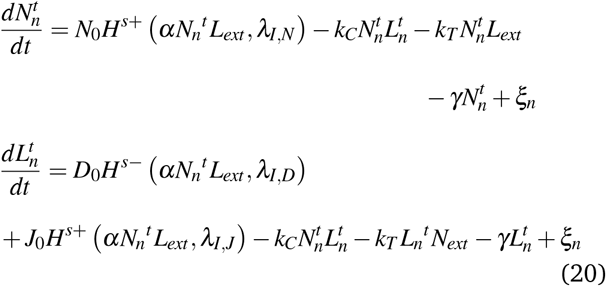

We expand the first term (the shifted Hill function) in Taylor series of Eq. (20) around the dissociation constant, *I*_0_. Considering the first two terms in the Taylor series expansion, we can further write Eq. (20) approximately as

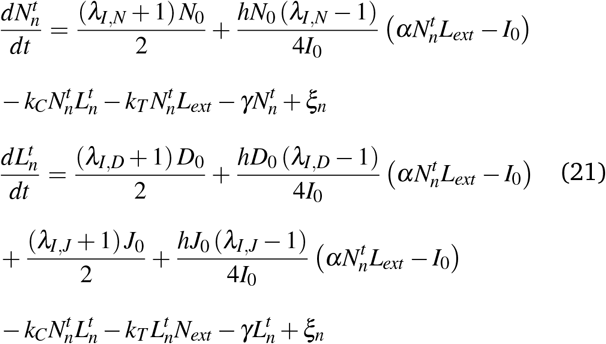

or in the matrix form, the above system of equations can be written as

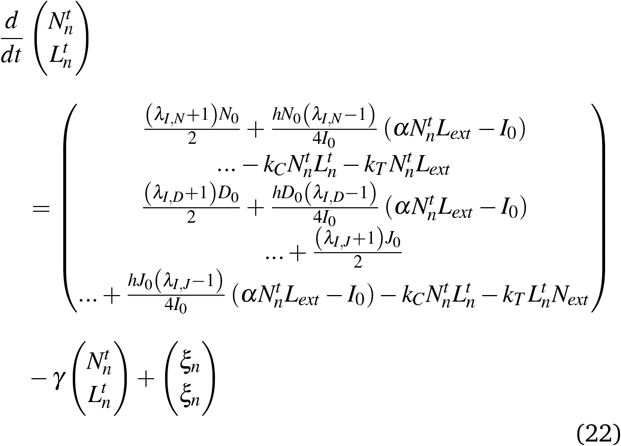

We absorb *γ* into the characteristic time scale of the system.We can now define the internal states as

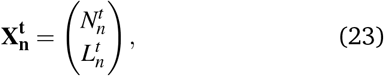

and external states as

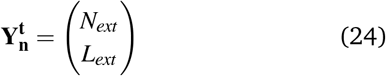

Using this definition of internal and external variables, we can write the system of Eq.(22) as

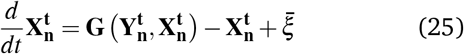

where

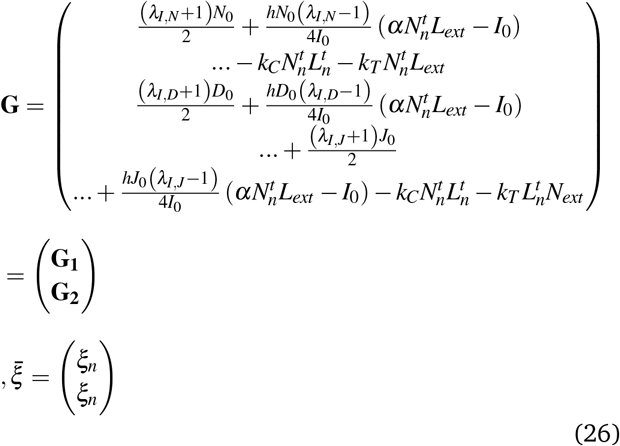

To find the steady state condition, we take 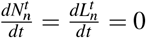. Now, we can use the tools previously mentioned in Eqs.(9) as

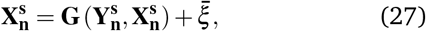

As we have defined the drift term and stead state equation, we are able to write the magnitude of the sensitivity parameter in the terms of drift as

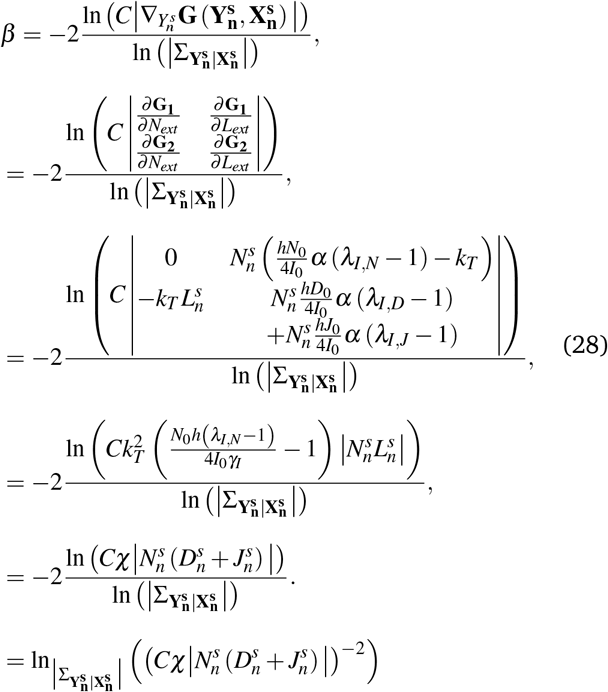

Here 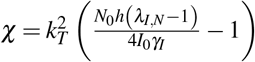. Please note that the probability distribution of other receptors present in the cell follows a Gaussian distribution. From the relation of Eq. (28), the value of sensitivity depends on the logarithm of internal variables and parameters, which are further modified by *Cχ*. From the above relation one can also find the condition of sensitivity whether it is 0, positive, or negative.

## 3 Results

### 3.1 Pattern formations

In this section, we have studied the pattern formation and the corresponding parameter regimes of the LEUP driven phenotypic mean-field model. From our simulations, it is seen that the four broad parameter regimes (as defined in Fig.1(B)) can be distinguished to study the pattern formation: there exists a clear difference between the nearest neighbor (*r* = 1) and extended neighbor (*r* > 1) interactions. Interestingly, we find that all extended neighborhood systems, from (*r* = 2) till, (*r* = 10) are qualitatively similar. The population distributions of phenotype change with differing values of sensitivity and radius of interaction, and these changes are classified into phase transitions 1 and 2. In the next sections, we shall discuss briefly the local variance and its particular role in phase transition regimes.

### 3.2 Order parameters to study phase transition regimes

For a qualitative description of phase transitions, we have to define an order parameter. In this particular scenario, we have chosen Sarle’s Bimodality Coefficient or the Hartigan’s Dip Statistic ^22^ and local variance as the order parameter. Hartigan’s dip test helps us to understand the multimodality in the sample points. On the other hand, as our model is based on entropy-based decision-making, and the conditional probability distributions were previously taken from Gaussian distribution for which entropy is a function of variance as 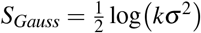), the measure of the overall entropy of the system could be defined in terms of local variances. For the *n* − *th* cell with phenotype 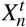, the arithmetic variance can be written as

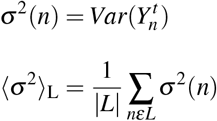

where *L* refers to the lattice and |*L*| is the number of lattice points. Thus, ⟨*σ*^2^⟩_L_ is the local variance, averaged over the entire lattice. Using the notion of local variance, one can understand the phase transitions and the phase diagram. Furthermore, its qualitative behavior and time evolution help us to identify a third phase transition (i.e., PT3)

### 3.3 Phase transition regimes

To understand the phase transitions 1 and 2 (PT1 and PT2), we have normalized the phenotypes of all cells between, [−1, 1] and their KDE plots are obtained for different conditions of (*β, r*). The two extrema *x* = 1 and *x* = −1 can be considered as two distinct phenotypes with intermediate values referring to hybrids.

#### 3.3.1 PT1

PT1 is observed for (*r* = 1) systems (considering only nearest neighbor interactions). **Fig.2(a)** and **Fig.2(b)** show *PT*1 in SQUARE and HEXAGONAL lattices respectively: populations go from being bimodal(unimodal) to trimodal(Cauchy or Lorentz-like distribution). In both cases, as *β* increases, the peak at *x* = 0 grows. This can be interpreted as: **Increased sensitivity, ⟹ increased frequency of hybrid phenotypes**. In square lattices, the cells seem to be driving their nearest neighbors to the phenotype/state opposite to their own (“lateral inhibition“). However, on the other hand in the case of hexagonal lattices: cells attempt to do the same due to the geometrical structure, the net signaling cancels out instead of adding up constructively, i.e., the “frustrated” systems. It could be that there exists no bimodality in hexagonal lattice populations. However, the population frequency of *x* = 0 (hybrid state) grows with increasing *β*, causing the KDE plots to go from being generically unimodal to Cauchy-Lorentz-like distributions with a pronounced peak at *x* = 0, consistent with what we define as *PT*1.

**Fig. 2.**
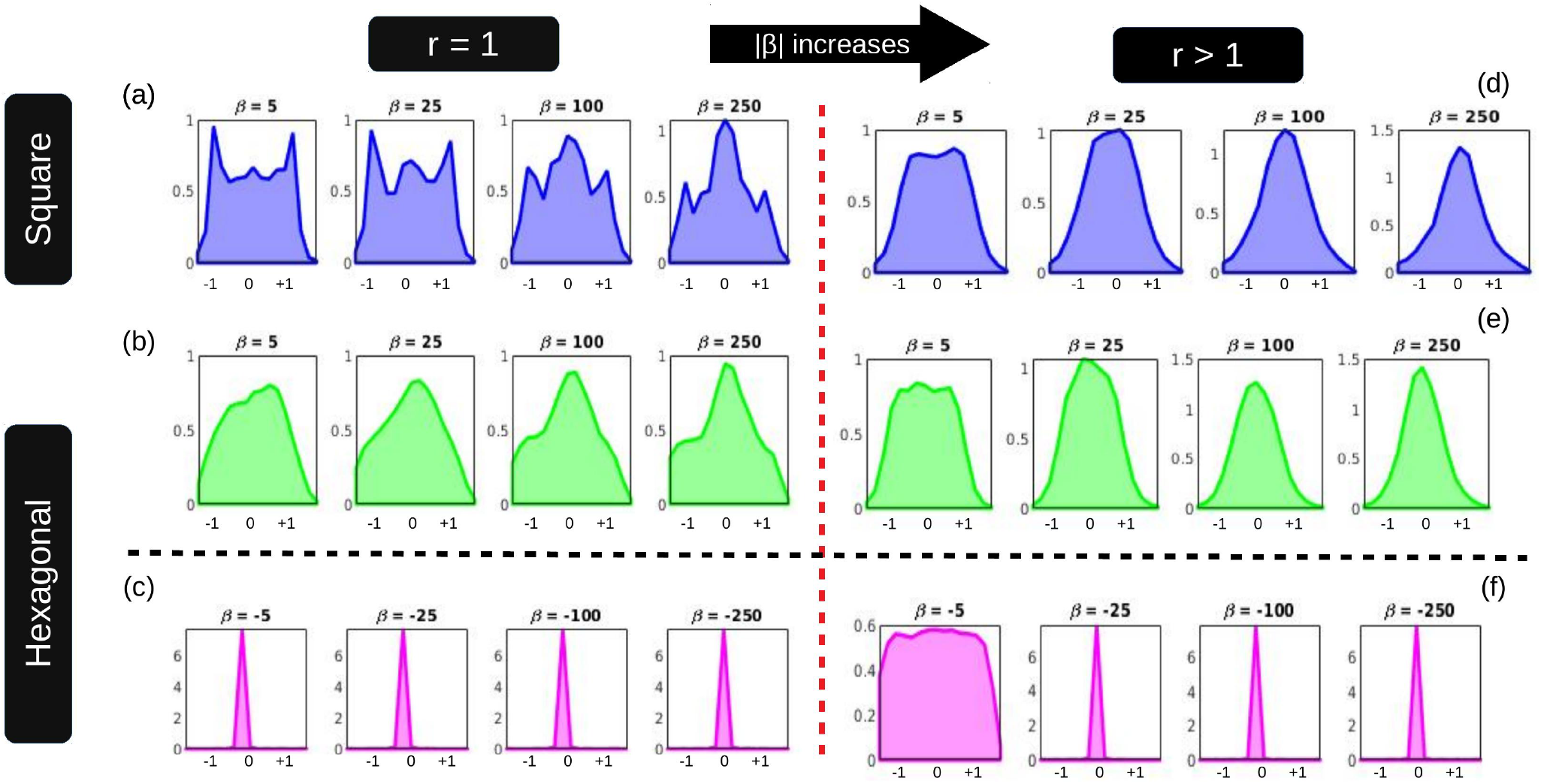
The steady state population distributions of phenotype exhibiting different phase transitions. The dotted black line separates plots for positive and negative, *β* while the dotted red line does so for (*r* = 1) and (*r* > 1). **(a)** and **(c)** Show *PT*1. **(e)** There is no change in the population distributions, thus for (*r* = 1, *β <*0), no PTs observed. All (*r* > 1) systems: **(b)**, **(d)** and **(f)** show *PT*2, albeit in slightly different ways.

#### 3.3.2 PT2

As seen in **Fig.2(b), Fig.2(d)** and **Fig.2(f)**, *PT*2 is observed in all (*r* > 1) systems, regardless of *β*. In all cases, for small to intermediate values of |*β*|, the populations are uniformly randomly distributed. This means the populations are merely evolving stochastically. As |*β*| increases beyond some threshold value, the sensitivity is enough for the cells to take informed decisions. For positive *β*, this leads to unimodal populations centered around *x* = 0 (the hybrids) with some amount of variance, while for negative *β*, almost all cells are driven to *x* = 0 phenotype, with few outliers, leading to *δ*-function like distributions. Phase Transition 2 is defined as the point beyond which populations show some patterning instead of remaining randomly distributed.

Moreover, as *r* increases, as we are considering contribution from a larger number of cells, the net signaling cancels out and grows weaker. ^*^ Accordingly, the threshold value of |*β*| required for appreciable sensing also increases.

The interpretation could be that: **Increased radius of interaction**, ⟹ **net signaling attenuates**. Thus, as the phase transitions are defined based on histograms of the phenotype distribution, indices that characterize unimodal or multimodal probability distributions such as the Sarle’s Bimodality Coefficient or the Hartigan’s Dip Statistic could be considered as order parameters. This is subject to the limitations of the indices themselves, as they may not always accurately classify a distribution. (Check S.I. for phase diagrams).

#### 3.3.3 PT3

PT3 is only observed in SQUARE lattices for (*r* = 1*, β* > 0) systems (in **Fig.3(A)i**) for very small values of *β*, ⟨*σ*^2^⟩_L_(*t*) decreases with time. For intermediate to higher values, it first shows a sharp decrease before increasing and then plateauing. The final values ⟨*σ*^2^⟩_L_ saturate with respect to (do not follow a monotonic trend) *β* - initially there is an increase as *β* increases, a change we have classified as *PT*3. (Changes upon further increasing *β* are indicative of *PT*1 occurring.). For all other parameter regimes, ⟨*σ*^2^⟩_L_ is a monotonically decreasing function of time. For any value of *β*, as *r* increases, the ⟨*σ*^2^⟩_L_ time evolution comes to resemble that of populations evolving stochastically (decrease is less defined), indicative of *PT* 2 . Trends in other order parameters are also consistent with the changes occurring in *PT*2 as well as *PT*1. To better understand how ⟨*σ*^2^⟩_L_ time evolution changes with *β*, the average slopes over all time, i.e. 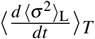 for different sensitivities are plotted against *β* in **Fig.3(B)**. A population that was evolving in an uninformed way would have 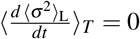, as at each point in time, the phenotypes would be uniformly randomly distributed and so would remain at the same values of ⟨*σ*^2^⟩_L_. Next, we shall study the phase transition regimes for different values of interaction radius.

**Fig. 3.**
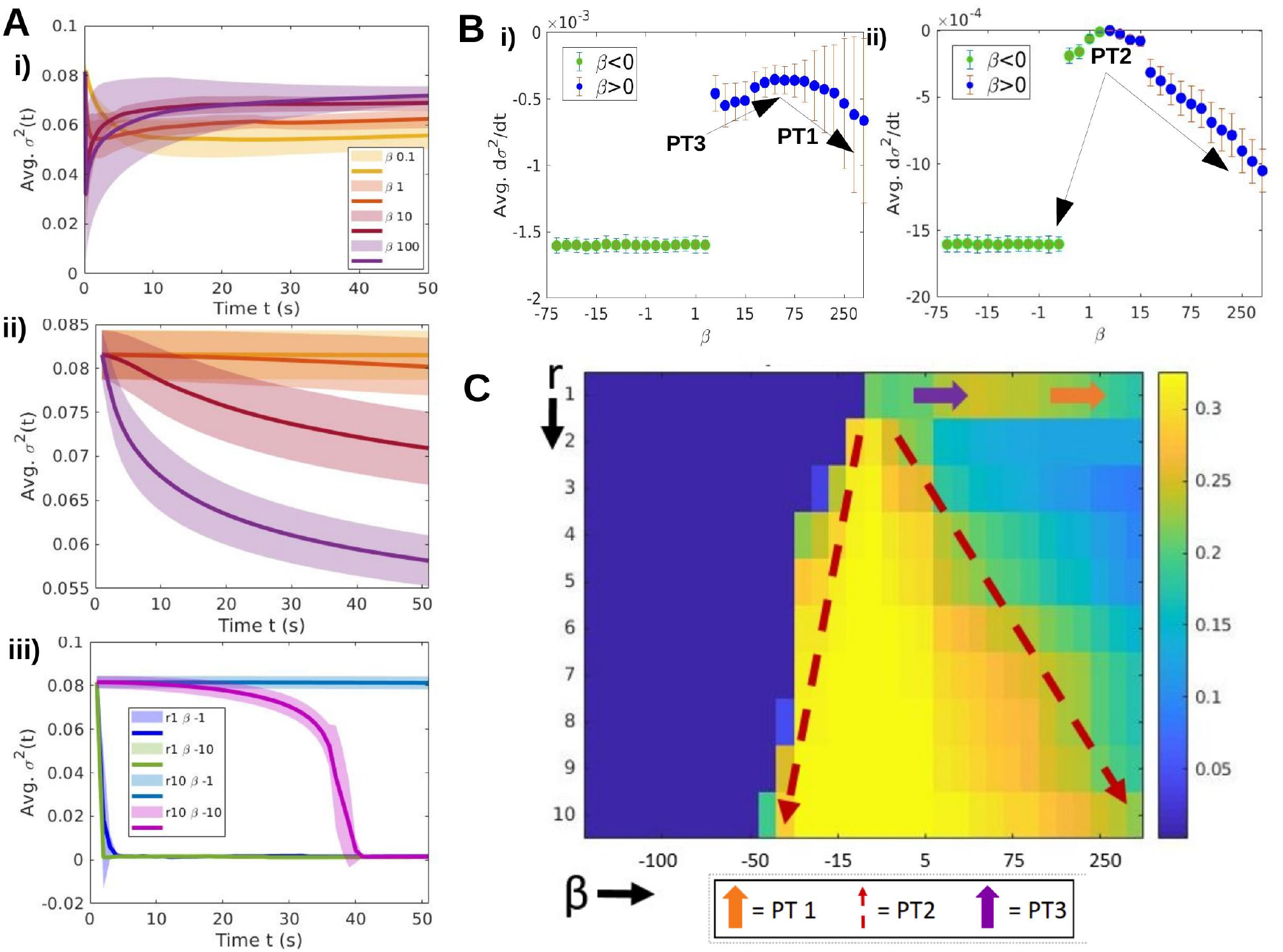
**(A)** Time evolution of local variance in square lattices. The x-axis is time in (100s). **(i)** (*r* = 1, *β* > 0): *PT*3 is observed; as *β* increases, ⟨*σ*^2^⟩_L_ goes from decreasing to increasing with time **(ii)** (*r* = 5, *β* > 0) and **(iii)** (*β* < 0) show monotonic trends. The local variances decrease with time. **(B)** The average slope of the time evolution graphs are plotted against *β* values for **(i)** *r* = 1 and **(ii)** *r* = 5. **(C)** Phase diagram with steady state ⟨*σ*^2^⟩_L_ as the order parameter.

**Table 1.**
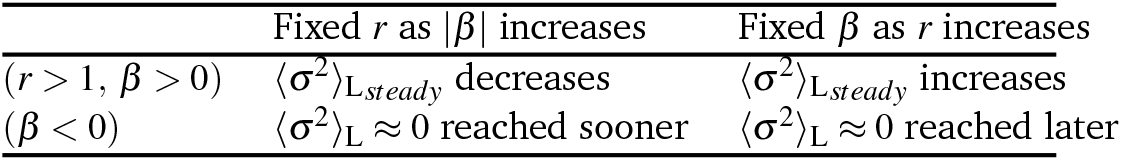
Qualitative behavior(s) observed when ⟨*σ*^2^⟩_L_ decreases with time

### 3.4 Phase transitions with different interaction radius

We have also found significant changes in PT1, PT2, and PT3 phase transition regimes with different interaction radius cases. We ran our simulations from small interaction radius to large interaction radius while fixing the sensitivity parameter i.e., |*β*|.

#### 3.4.1 Small interaction radius case (r = 1 system)

For negative values of *β*, no phase transition is observed for (r=1) system as expected. Even very small values of |*β*| (=0.1) are enough for sensing and cells are driven to the *δ*-function like distributions centered around *x* = 0. On the other hand, for positive values of *β*, we have already seen that the initial increase in 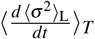 can be attributed to *PT*3. Upon further increasing *β*, there is a decrease in both, 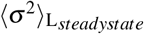 and time-averaged slope. This can be attributed to *PT*1 taking place (consistent with the fact that trimodal populations (*β* = 250) have a lower variance than bimodal populations (*β* = 25)).

#### 3.4.2 Large interaction radius case (r = 5 system)

In case of both positive and negative *β*, *PT*2 is hinted for (r=5) system. There is not much sensing for small, |*β*| so the average slope values are close to 0. Beyond |*β*_*critical*_|, when *PT*2 occurs and the cells start taking informed decisions, there is a sudden decrease in the average slope. The shift seems more abrupt or “discontinuous” in case of negative *β*, while it seems more gradual for positive sensitivity.

#### 3.4.3 Phase diagram

**Fig3(C)** is a phase diagram with 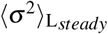 as the order parameter for square lattices. *PT* 3 and *PT* 1 as well as the clear distinction between (*r* = 1) and (*r* > 1) systems is seen. For each value of *r* along the dotted red lines labelled *PT* 3, the phase transition is occurring for different critical values of *β*. As previously mentioned, |*β*_*c*_| is increasing with *r* for positive as well as negative values of *β*. Thus, **Fig3(C)** Summarizes all the types of patterns seen in the cell populations and using the order parameter ⟨*σ*^2^⟩_L_, we are now in a position to draw qualitative parallels between the systems governed by LEUP model to the patterning observed in the celebrated biological system of Notch-Delta signaling. If the system is Delta dominated, i.e, *D*_0_ is large, then we see salt-pepper patterning. In the square lattice, we get check-board type of patterns, the cells choosing either of the two possible states. In the hexagonal lattice, we see more interesting patterns due to frustration in the lattice. One possibility is that the cell at the center chooses the high Notch-low Delta state, whereas the surrounding cells in the hexagon, chose the opposite fate. The vice versa is also possible. This results in an increase in *σ*^2^ over time. If Jagged signaling is dominant, i.e., *J*_0_ is large, the system goes to a uniform phenotype. All cells tend to attain the same phenotype. This results in *σ*^2^ decreasing over time.

Notch Delta signaling occurs predominantly through receptors and ligands attached to the surface of cells, making it a strictly nearest-neighbour interaction, i.e. there is no equivalent in the Notch Delta systems for *r* > 1 systems. Thus, parallels are only drawn between them and (*r* = 1) LEUP populations.

### 3.5 Comparison between mechanistic model and entropy-based model

In this section, we compare our results of entropy based model with the classical Notch-Delta-Jagged model. For the resemblance, we use two kinds of lattice setups i.e., (**1**) 1–17 | 11 square lattice and (**2**) hexagonal lattice. The time evolution of average local variance^†^, ⟨*σ*^2^⟩_L_ helps us to draw qualitative parallels between the two models, which are further quantified with the help of the radial distribution function. There are two overarching cases.

#### 3.5.1 Lateral Induction: (Decreasing ⟨*σ*^2^⟩_L_)

These are compared in **Fig.(4)**. In LEUP populations with *β* < 0, ⟨*σ*^2^⟩_L_ decreases with time and all the cells approach the same phenotype *x* = 0. The same uniform patterning as well as time evolution behaviour of ⟨*σ*^2^⟩_L_ (decreasing with time) is observed in Jagged-signalling dominated NDJ systems. In both square and hexagonal lattices, the LEUP and NDJ models compare very well for this parameter regime. It is notable that while NDJ systems give rise to perfect lattices in which all cells are driven to the same phenotypes, LEUP populations always have some outliers/imperfections in the lattice^‡^

**Fig. 4.**
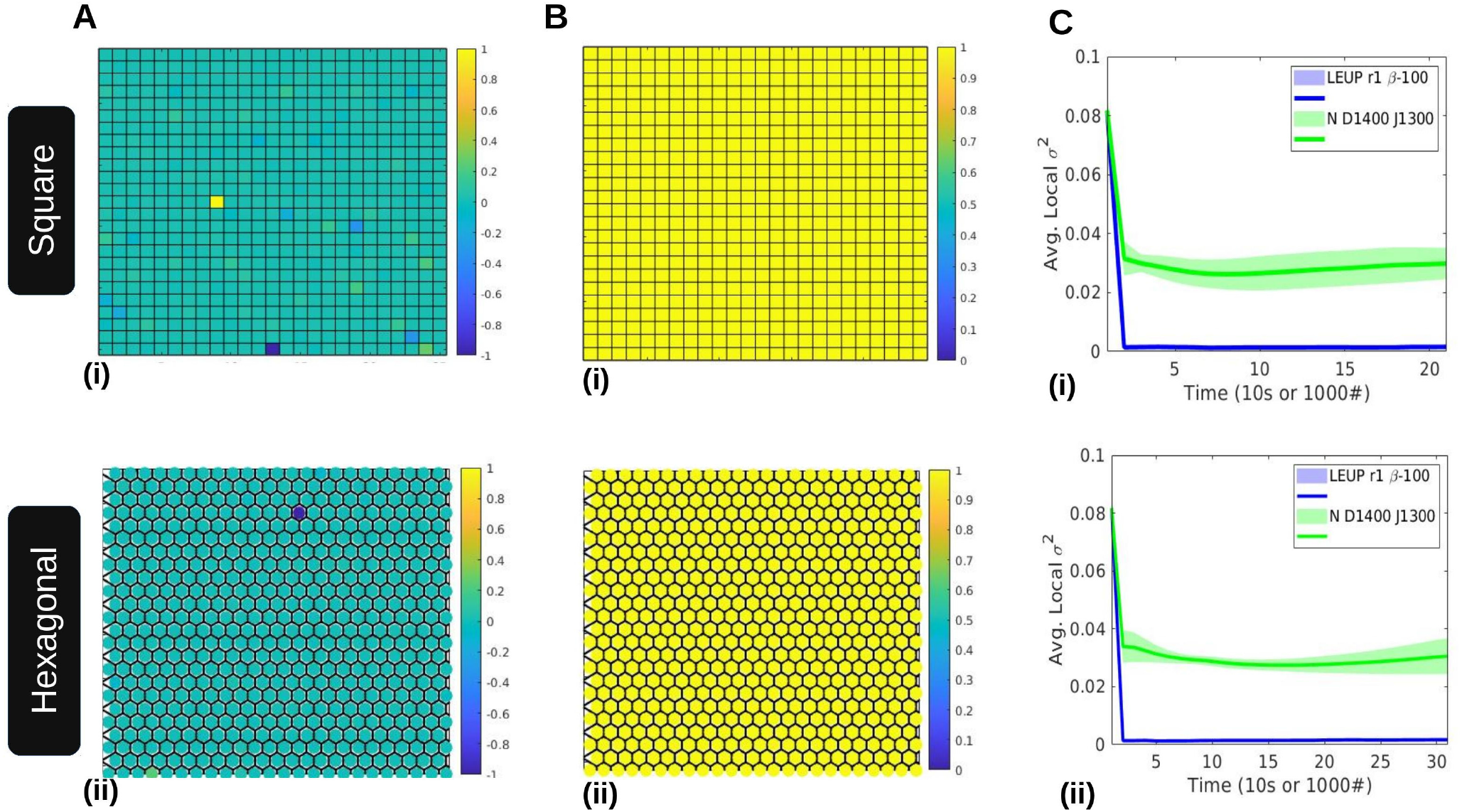
Spatial patterning seen in **(A)** LEUP **(B)** Notch Delta systems are visualised (Heatmaps plotted of phenotype and NICD levels respectively). **(C)** ⟨*σ*^2^⟩_L_ decreases with time in both cases. (Plots are averaged over 10 simulations and the shading is representative of the standard deviation in them)

#### 3.5.2 Lateral Inhibition: (Increasing ⟨*σ*^2^⟩_L_)

This case is studied in **Fig.(5)**

**Fig. 5.**
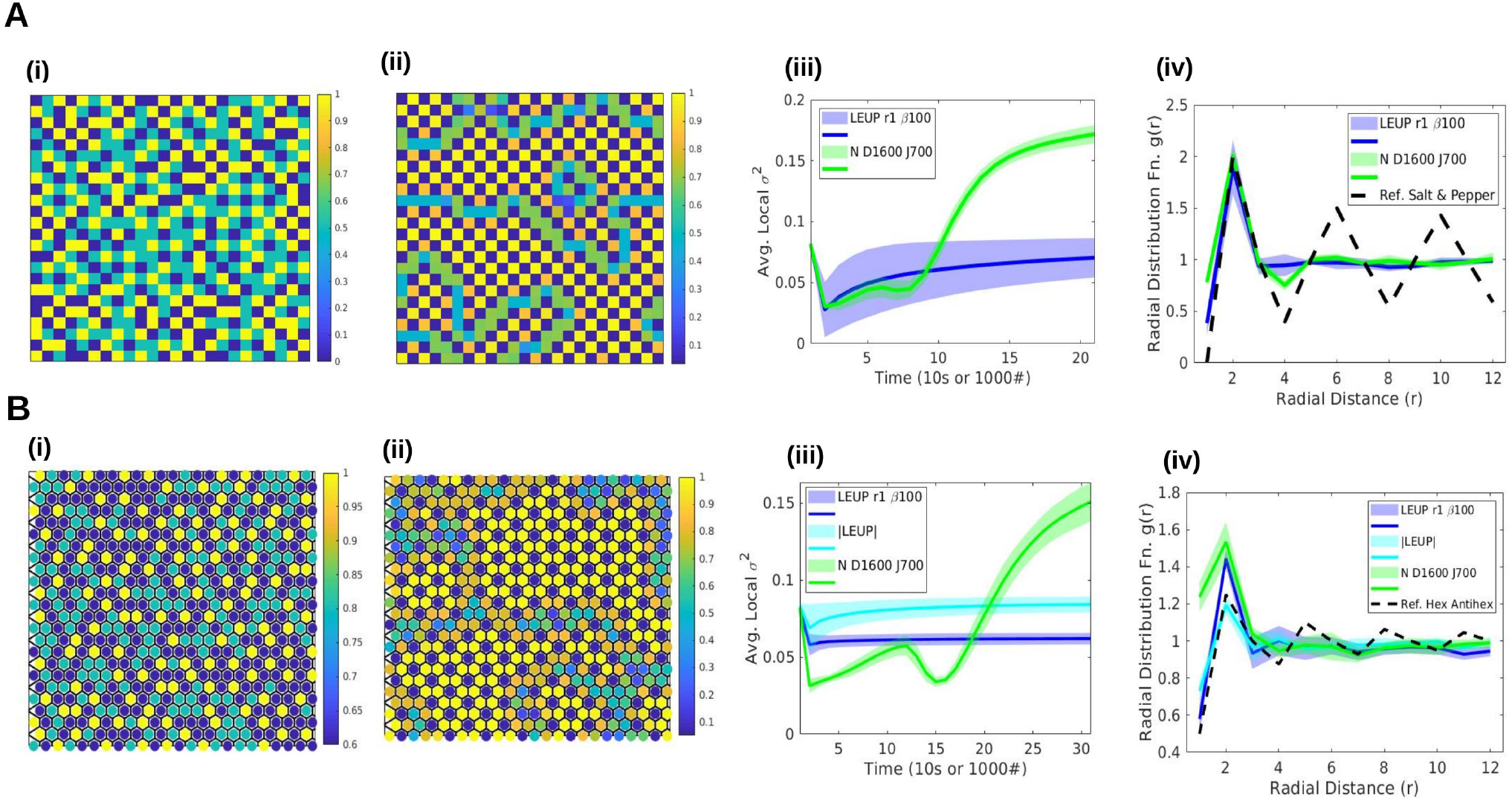
**(A)** Square Lattice. Heatplots of *(i)* phenotype in LEUP(*r* = 1, *β* = 100) and *(ii)* NICD levels in NDJ(*D*_0_ = 1600, *J*0 = 700) systems. To make the patterns more evident, the continuous variables have been discretised. *(iii)* shows time evolution of ⟨*σ*^2^⟩_L_. *(iv)* Radial distribution function, *g*(*r*). **(B)** Hexagonal Lattice. While *(ii)* is the heatmap of NICD levels in the NDJ system, *(i)* is the discretised heatmap of *absolute values of phenotype*, |*x*|. The *(iii)* ⟨*σ*^2^⟩_L_ time evolution graphs and *(iv)* radial distribution functions contain plots of both, *LEUP* and |*LEUP*|.

##### Square Lattice

The square lattice cases compare quite well. A distinct chessboard-like (“salt and pepper“) patterning is observed in patches in both cases, although there is significantly higher order in the NDJ system. The black dotted line in the radial distribution function plots is of a reference (perfect) chessboard pattern to quantify the similarity in spatial patterning. The local variance time evolution plots are also similar- an initial decrease, subsequent increase, and then saturation in ⟨*σ*^2^⟩_L_ values. However, the two systems seem to be on different time scales - LEUP has a much sharper decrease and saturates more quickly than the NDJ. Due to frustration in the boundaries of salt pepper patterns, one can observe “fluid”-like arrangements in the phenotypic space from the radial distribution function in both models.

##### Hexagonal Lattice

When trying to directly compare, the parallels between hexagonal systems fall short. The NDJ system adopts a “hexagon-antihexagon” type of patterning. (In the radial distribution function plots, **Fig.5B(iv)**, the dotted line is of the same reference pattern). This is not replicated by the LEUP simulations. Moreover, the local variance is increasing in time for the former and decreasing for the latter. However, the LEUP populations do show a less obvious pattern - cells at either extrema, *x* = ±1 are mostly surrounded by hybrid phenotypes, *x* = 0. Thus, upon taking the absolute values of phenotypes as the order parameter instead, the spatial patterns show some degree of resemblance to those of the Notch Delta simulations. The local variance of the absolute values of phenotype is also increasing with time (cyan plot, Fig.5B(iii)). Similar to square lattice, one can inspect the “fluid”-like trend in the phenotypic space for the hexagonal lattice.

### 3.6 Other spatial patterns

The one class of distributions whose spatial patterning we have not discussed so far are the (unimodal) *LEUP*(*r* > 1, *β* > 0) systems. Here too, cells of extreme phenotypes are almost always surrounded by cells of hybrid (*x* = 0) phenotypes; very rarely are more than 2-3 cells of *x* = ±1 immediate neighbors. While this patterning is not easily defined or immediately obvious to the human eye, it is captured in the spatial Fourier transforms of the heatmaps: while checkerboard-like distributions have FFTs localized at the center, the FFT of these unimodal populations has an almost ring-like structure (**Fig.6B**). This is further verified by the radial power spectra (**Fig.6C**) in which the peak of the unimodal populations is at a higher radial distance from the center. Though the concept of extended neighborhoods is not directly applicable to NDJ populations, the above patterning has some similarities with those observed in early patterning in the pancreas, in which the Notch Delta signaling adopts a “lateral stabilization” mechanism.

**Fig. 6.**
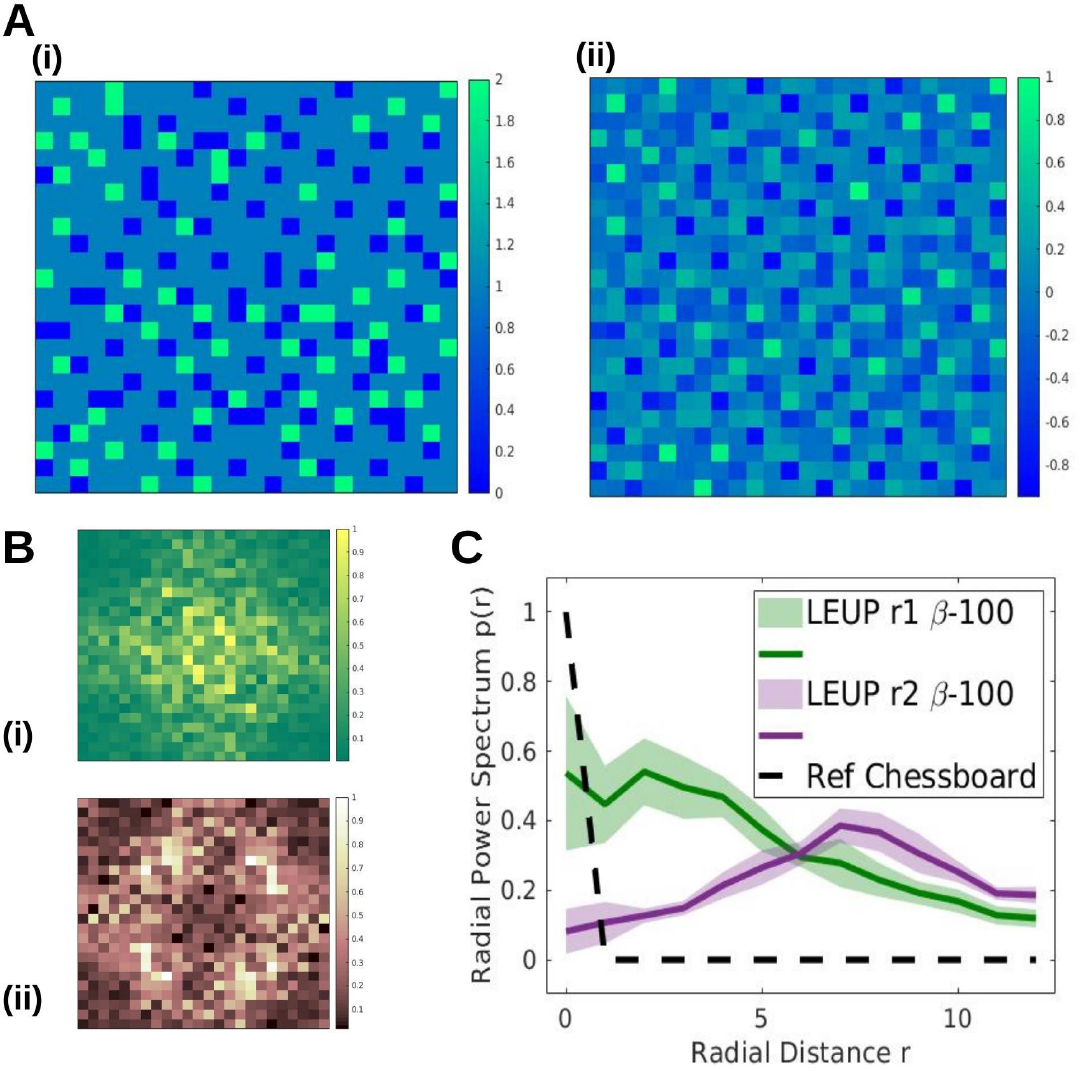
**A** Heatmaps of LEUP(*r* = 2, *β* = 25) in a square lattice, of **(i)** discretized and **(ii)** continuous phenotypes. Isolated ‘islands’ of extrema cells surrounded by hybrid phenotypes are evident. **B** Magnitudes of spatial Fourier transforms of **(i) “**salt and pepper” patterned LEUP(*r* = 1, *β* = 25) and **(ii)** LEUP(*r* = 2, *β* = 25) populations. **(C)** Radial power spectrum of the spatial Fourier transforms of the two populations and a reference chessboard pattern.

### 3.7 Calculating *β* from NDJ simulations

We seek to further corroborate our numerical/simulations based understanding of the role of *β*, the sensitivity parameter with the analytical values we obtain from Notch Delta Jagged simulations. From (30) we have

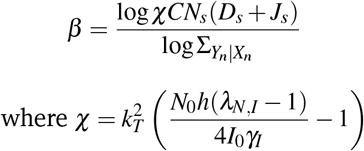

**Table 2.**
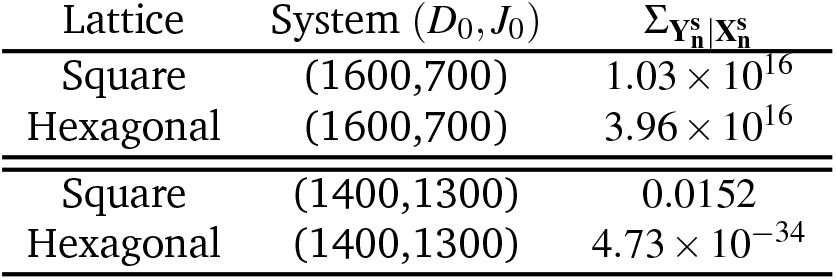
Calculating Covariance Matrix Determinants

The steady states levels of the receptors and ligands are first obtained from 1-cell and2-cell simulations and then compared with the values obtained for lattices. (This is in order to ensure no errors due to finite size effects of the lattice have not crept in). A ballpark estimate of the Σ_**Y**|**X**_, we calculated it from the lattice based simulations thus: having defined 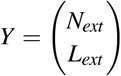 and 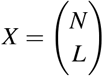, For multivariate Gaussian distributions,

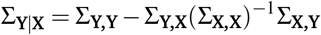

Here, we have

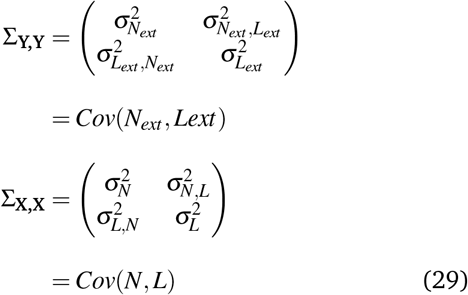

and

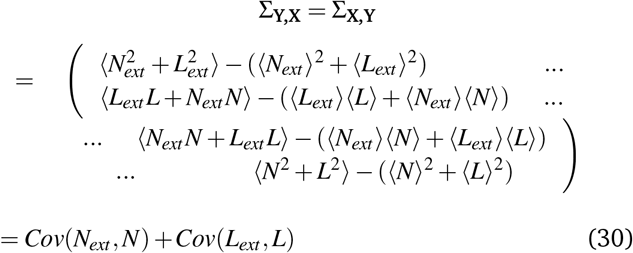

The values of *β* are calculated and tabulated in Table.(3). Due to our assumptions, though the values of *β* are not exact, it is striking that the signs come out to be the same as those obtained in the agent-based lattice simulations.

**Table 3.**
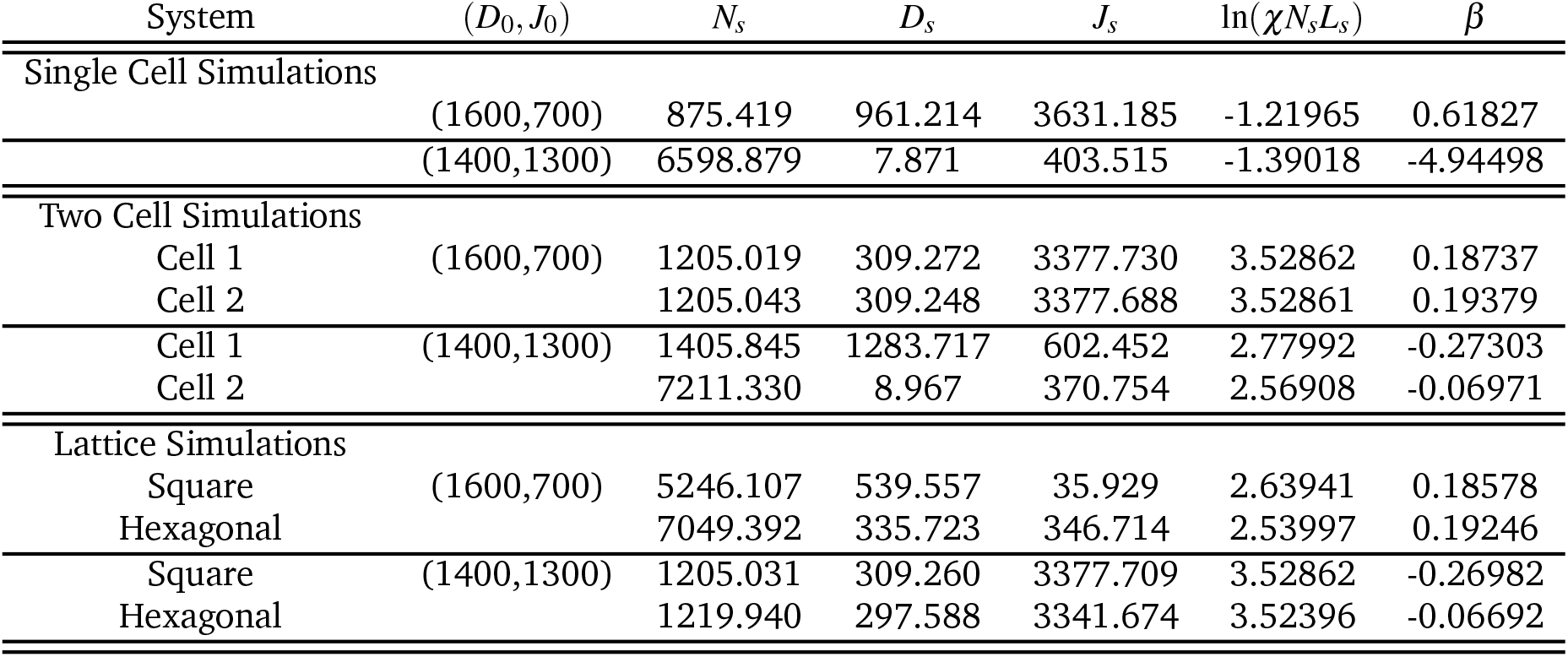
Calculating *β*

| System | $(D_0, J_0)$ | $N_s$ | $D_s$ | $J_s$ | $\ln(\chi N_s L_s)$ | $\beta$ |
| --- | --- | --- | --- | --- | --- | --- |
| Single Cell Simulations |  |  |  |  |  |  |
|  | (1600,700) | 875.419 | 961.214 | 3631.185 | -1.21965 | 0.61827 |
|  | (1400,1300) | 6598.879 | 7.871 | 403.515 | -1.39018 | -4.94498 |
| Two Cell Simulations |  |  |  |  |  |  |
| Cell 1 | (1600,700) | 1205.019 | 309.272 | 3377.730 | 3.52862 | 0.18737 |
| Cell 2 |  | 1205.043 | 309.248 | 3377.688 | 3.52861 | 0.19379 |
| Cell 1 | (1400,1300) | 1405.845 | 1283.717 | 602.452 | 2.77992 | -0.27303 |
| Cell 2 |  | 7211.330 | 8.967 | 370.754 | 2.56908 | -0.06971 |
| Lattice Simulations |  |  |  |  |  |  |
| Square | (1600,700) | 5246.107 | 539.557 | 35.929 | 2.63941 | 0.18578 |
| Hexagonal |  | 7049.392 | 335.723 | 346.714 | 2.53997 | 0.19246 |
| Square | (1400,1300) | 1205.031 | 309.260 | 3377.709 | 3.52862 | -0.26982 |
| Hexagonal |  | 1219.940 | 297.588 | 3341.674 | 3.52396 | -0.06692 |

Lateral Inhibhition (*D*_0_, *J*_0_) = (1600, 700) ⟹ *β* > 0

Lateral Induction (*D*_0_, *J*_0_) = (1400, 1300) ⟹ *β* < 0

## 4 Conclusions

In this article, we have proposed a mathematical model of lattice-based phenotypic decision-making. We assumed that each cell in the lattice will change their phenotype according to the entropy of their microenvironment.

We exhibited the origin of sensitivity parameter. We found that in the low noise approximation limit, the sensitivity parameter is the ratio of logarithmic factors of (**I**) the gradient of the drift and (**II**) the variance of the microenvironmental distribution. Furthermore, we use the example of the Notch-Delta-Jagged signaling to estimate the sensitivity parameters. Using this example, we have established the conditions for which the sensitivity parameter can have three possible cases (i.e., *β* = 0, *β* < 0 and *β* > 0). In the agent-based model of phenotypic decisions, we characterize the pattern formation appeared from microenvironmental phenotypic interactions. Furthermore, we compared our model with classical mechanistic Notch-Delta-Jagged signaling. To study this, we use two neighborhoods i.e., (**a**) the nearest neighborhood (*r* = 1) and (**b**) the extended neighborhood (*r* > 1) interactions. For the nearest neighborhood interactions, the distribution of phenotypes are changing from bimodal to trimodal distribution for both lattices (i.e., square and hexagon) when we increase the sensitivity parameter. In the square lattice, we found salt-pepper patterns which resembled the patterns found in the mechanistic Notch-Delta-Jagged model, and in the hexagonal lattice we could not find the signatures of bimodality due to the frustration of taking decisions. Interestingly, for the extended neighborhood (*r* > 1) interactions, we found unimodal distributions as we escalate the sensitivity parameter values for both lattices. Additionally, we used the radial distribution functions and average local variance to quantify the phenotypes over space and explored the “fluid”-like structures developed in the phenotypic space for both square and hexagonal lattices.

Notch signaling, an evolutionary conserved intercellular communication pathway, coordinates the emergent spatiotemporal patterning in a tissue via interactions between Notch receptors and different transmembrane ligands (Delta, Jagged etc.) of neighboring cells and regulates cell-fate decisions. This pathway is ubiquitous in nature and is responsible for various biological processes, viz., embryonic development, wound healing, angiogenesis, and cancer progression ^23^. Intracellular interplay between Notch-Delta-Jagged signaling and transcription factors and microRNAs based regulatory network for Epithelial Mesenchymal Transition (EMT) ^24^ during cancer metastasis might result in other different interesting spatiotemporal patterns at tissue level. This Notch-EMT combined mechanistic circuit would have a different microenvironment. So, developing an LEUP framework for this scenario could provide more insights for the spatiotemporal patterning in cancer metastasis, in particular.

The underlying causes for phenotypic changes in a cell are hard to know. To model this, we use the information from the microenvironment in the form of entropy, which will further guide the individual cells. Here the cell will follow the LEUP and try to minimize the uncertainty over time. Although, we assumed that the distribution of the microenvironment is taken from Gaussian distribution, one can use a different microenvironmental distribution to see the underlying patterns emerged from phenotypic decisions. If there exists a strong coupling between the microenvironment and the cell, one may use other definitions of entropy like non-ergodic entropy (e.g., Tsallis entropy ^25^, Renyi entropy ^26^ etc.), which will further help us to investigate phenotypic decision-making in a deeper level. In the future, we shall also consider the off-lattice case of phenotypic decision-making. We have not considered the phenomena of cell proliferation here. It will be interesting to see how the rate of proliferation can influence the pattern formation regimes. Moreover, one can study the phenotypic decision-making in different topologies (from networks to lattices and spatial geometries). We have used periodic boundary conditions in the simulations. It will be nice to explore the impacts of other different boundary conditions (e.g., reflective and absorbing boundary conditions etc.) on phenotypic decision-making on the spatial scale.

In summary, we have shown how the microenvironmental uncertainty can drive the phenotypic decision-making on on-lattice, which further help us to study phenotypic decision-making when the precise knowledge about internal states of the cells are unknown and/or partially known.

## Acknowledgements

A.P. and D.S. are supported by KVPY fellowship awarded by Department of Science and Technology (DST), Government of India. UR acknowledges C. V. Raman Postdoctoral Fellowship from Indian Institute of Science, Bangalore, India. MKJ is supported by Ramanujan Fellowship (SB/S2/RJN 049/2018) awarded by the Science and Engineering Research Board (SERB), DST, Government of India. A.B. and H.H. would like to acknowledge the VolkswagenStiftung funding within the *Life?* program (96732). H.H. is supported by MiEDGE (01ZX1308D) of the ERACOSYSMED initiative. Moreover, H.H. acknowledges the support of the FSU grant 2021-2023 grant from Khalifa University. A.B. 1–17| 15 and H.H. also thank the Centre for Information Services and High-Performance Computing (ZIH) at TU Dresden for providing an excellent infrastructure.

## 5 Supplementary Material

### 5.1 Estimating the partition function Z^′^

By definition, we have

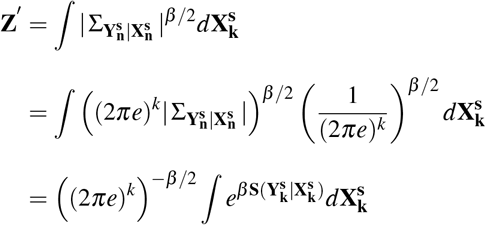

From the identity *S*(**X**, **Y**) = *S*(**X**) + *S*(**Y**|**X**) ≤ *S*(**X**) + *S*(**Y**), we know *S*_*max*_(**Y**|**X**) = *S*(**Y**).

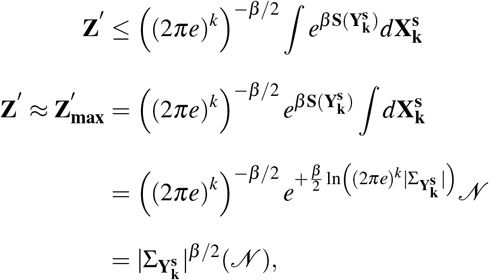

where 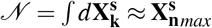, a biologically set upper limit on the number of receptor molecules.

### 5.2 Effect of different neighborhoods on *β*_*critical*_ of *PT*2

From Fig.(7), we can firstly see that for all neighborhoods, with increasing radius of interaction or number of neighbours considered, the *β*_*critical*_ for *PT*2 seems to almost monotonically increase. This further underlines our inference that too great a neighborhood of interaction leads to net signal attenuation.

**Fig. 7.**
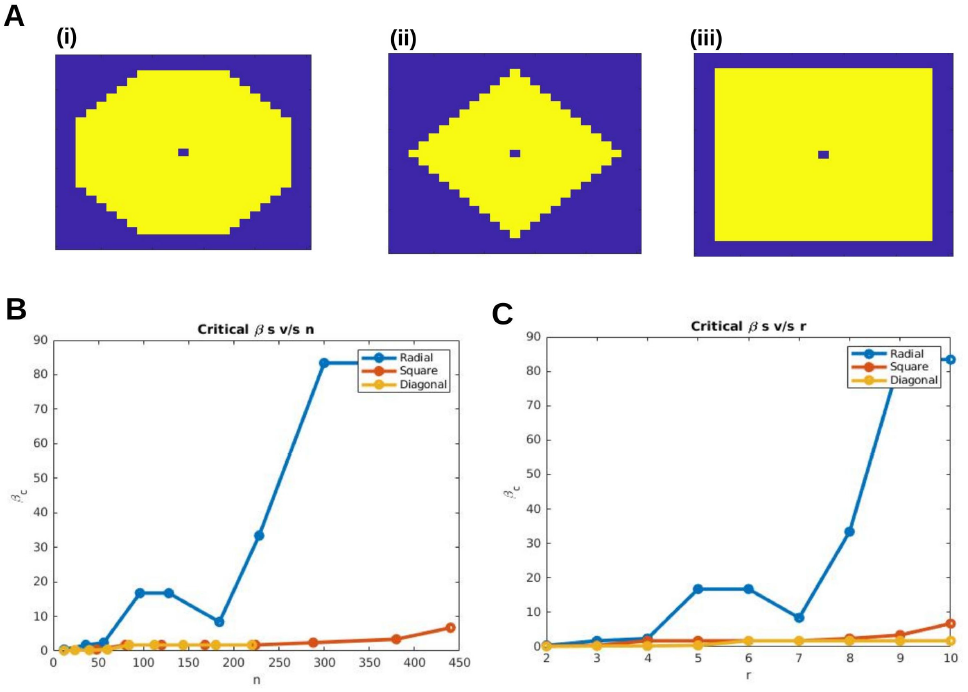
**(A)** Shows the three different kinds of neighborhoods considered in square lattices, **(i)** radial, **(ii)** diagonal and **(iii)** square. **(B)** *β*_*critical*_ as a function of *r* and *n* (number of neighbours) for the different neighborhoods

### 5.3 Phase Diagrams using Statistical Measures as Order Parameters

The Sarle’s Bimodality Index enables us to differentiate between unimodal (BC*<*0.555), multimodal (BC*>*0.555) and uniformly distributed (BC=0.555) populations. It is therefore a useful tool to quantify the critical parameter values for the observed phase transitions and their nature is hinted at in phase diagrams using this index as an order parameter. (Fig.(8))

**Fig. 8.**
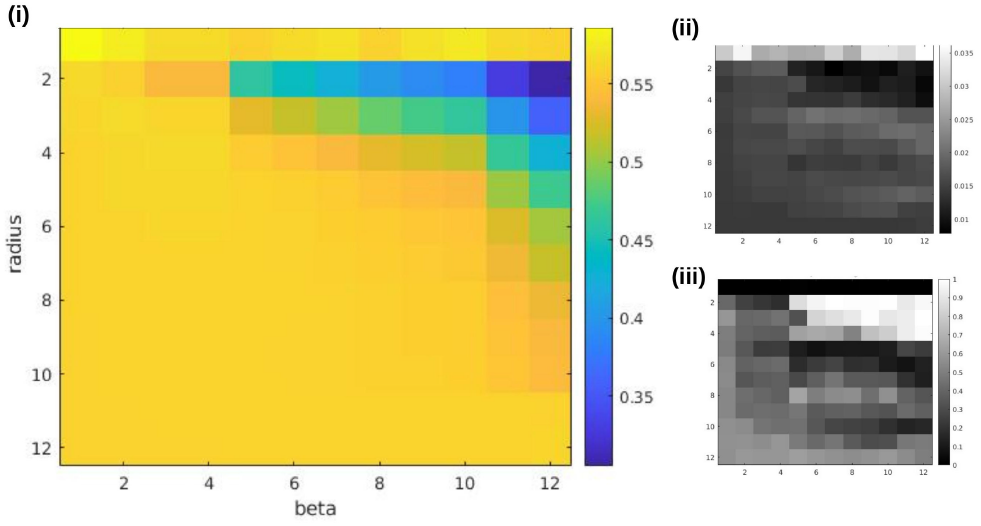
Phase diagrams of **(i)** Sarle’s Bimodality Coefficient **(ii)** Hartigan’s Dip Statistic **(iii)** p-value of significance test on the dip statistic with the null hypothesis being that the population is unimodal

## Footnotes

* For different types of neighborhoods, the number of cells in the neighborhood changes differently. S.I. contains further details

† When we say ⟨*σ*^2^⟩_L_ is increasing in time, we simply mean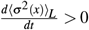 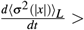,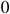, 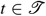, for some interval 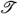. The populations will eventually reach steady state when ⟨*σ*^2^⟩_L_ no longer changes with time

‡ This is without incorporating any noise terms too, all simulations are deterministic.

